# Poly-basic peptides and polymers as new drug candidate against *Plasmodium falciparum*

**DOI:** 10.1101/2023.09.16.558069

**Authors:** Roshan Sivakumar, Katherine Floyd, Erath Jessey, Jenny Kim Kim, Peter O. Bayguinov, James A.J. Fitzpatrick, Dennis Goldfrab, Marko Jovanovic, Abhai Tripathi, Sergej Djuranovic, Slavica Pavlovic-Djuranovic

## Abstract

*Plasmodium falciparum*, the malaria-causing parasite, is a leading cause of infection-induced deaths worldwide. The preferred treatment approach is artemisinin-combination therapy, which couples fast-acting artemisinin derivatives with longer-acting drugs like lumefantrine, mefloquine, and amodiaquine. However, the urgency for new treatments has risen due to the parasite’s growing resistance to existing therapies. Our study shows that a common characteristic of the *P. falciparum* proteome – stretches of poly-lysine residues such as those found in proteins related to adhesion and pathogenicity – can serve as an effective peptide treatment for infected erythrocytes. A single dose of these poly-basic peptides can successfully diminish parasitemia in human erythrocytes *in vitro* with minimal toxicity. The effectiveness of the treatment correlates with the length of the poly-lysine peptide, with 30 lysine peptides supporting the eradication of erythrocytic parasites within 72 hours. PEG-ylation of the poly-lysine peptides or utilizing poly-lysine dendrimers and polymers further increases parasite clearance efficiency and bolsters the stability of these potential new therapeutics. Lastly, our affinity pull-downs and mass-spectrometry identify *P. falciparum’s* outer membrane proteins as likely targets for polybasic peptide medications. Since poly-lysine dendrimers are already FDA-approved for drug delivery, their adaptation as antimalarial drugs presents a promising new therapeutic strategy.

**One-Sentence Summary:** Our study demonstrates that poly-lysine peptides, particularly those modified through PEG-ylation or in the form of poly-lysine dendrimers, can effectively reduce *Plasmodium falciparum,* the causative agent of malaria, in human erythrocytes *in vitro,* with potential for use as a promising new antimalarial therapy.

## Introduction

In 2021, malaria affected an estimated 247 million people around the globe. This figure represents an increase from the 245 million cases reported in 2020 due to the COVID-19 pandemic (***1***). Despite a reduction in malaria case incidence (cases per 1000 population at risk) from 82 in 2000 to 57 in 2019, the incidence stayed the same or slightly increased in the following years, with 59 cases per 1000 in 2020 and 2021 (***1***). Human malaria, caused by *Plasmodium falciparum,* is responsible for over 600,000 deaths annually, mainly affecting children under age five. The *P. falciparum* life cycle includes two different hosts, humans and mosquitos. By implementing successful control measures such as the utilization of effective drugs for human treatment and efficient vector control strategies that target mosquitoes responsible for transmitting malaria, it is possible to achieve efficient control over infections and the spread of the disease. While malaria is less prevalent today than two decades ago, unfortunately, current drugs, including artemisinin and combination partners, are confronted with parasitic drug resistance (***2, 3***). As such, there is a need for new antimalarials that exploit novel mechanisms.

The current drug treatment paradigm for infected patients is artemisinin-combination therapy (ACT) (***4***). ACT is based on artemisinin derivatives, aggressive short-acting drugs, in combination with multiple long-acting drugs. Artemisinin gets activated by the cleavage of its endoperoxide, which then alkylates hundreds of proteins, potentially causing “proteinopathy,” eventually resulting in parasite death (***5, 6***). Different strategies were examined in the creation and further advancements of ACT or novel medications targeting malaria parasites. (***3, 7***). Some of these included the use of different combinations of existing drugs that might not be sufficient alone similar to ACTs, modification of existing drugs with promising candidates such as artefenomel and ferroquine being good candidate examples, and discovery of newer effective compounds (KAF156, amongst others) (***8***). However, it has also been shown that a range of promising antimalarial peptides, natural and synthetic, exhibit cytotoxic or inhibitory activity on malaria parasites. However, their clinical application has so far been limited (***9***). The most attractive feature of peptide-based therapeutics generally termed antimicrobial peptides (AMPs) (***9***), is their unique ability to affect multiple targets, thereby reducing the potential for resistance development by the targeted organism (***9, 10***). Finally, peptide-based drugs provide a superior platform for generating diverse medications by leveraging various biological and synthetic amino-acid analogs and modifications by non-amino-acid moieties (***9, 10***). The simple design, strong therapeutic outcomes, and their intrinsic malleability make anti-microbial peptides a promising avenue for exploratory drug discovery and development. Polycationic peptides are a member of the AMP family, which are lethal against many bacteria, fungi, and viruses (***11, 12***). The positive charge on the peptides allows them to interact with the negatively charged microbial cell membrane, leading to membrane disruption and cell death (***13***). This relatively non-specific mechanism of action can reduce the risk of resistance development compared to traditional antibiotics (***14***). Poly-L-lysine (PLL), a highly charged polycationic peptide, has been studied for several applications, including its antimicrobial potential (***15***).

*P. falciparum* has a highly AT-rich genome (81%) with coding sequences close to 75% AT-richness (***16***). Previous work has indicated that *Plasmodium* species are among the few rare organisms that can translate long runs of poly-adenosine in mRNA transcripts to poly-L-lysine peptides with high accuracy (***17***). Almost 50% of the *P. falciparum* genome codes for proteins with poly-L-lysine stretches, with some proteins containing 40 lysine residues in a row (***17–19***). In most organisms, poly-lysine, and more generally poly-basic residue-rich genes, are typically associated with RNA biogenesis, DNA repair, and chromosome segregation (***20***). However, the poly-lysine motif-containing proteins of *P. falciparum* and other *Plasmodium species* are also affiliated with adhesion and among the secretion repertoire of proteins involved in malaria pathogenesis and invasion (***21***). The hypothesis for this enrichment of positively charged amino acid patches in *Plasmodium* adhesion and pathogenesis is that parasites utilize polybasic loops in secreted and membrane-associated proteins to attach and invade host erythrocytes (***21–23***). This hypothesis is partially supported by previous studies in which negatively charged heparin or glycosaminoglycan blocked *P. falciparum* growth and invasion (***22, 23***). However, charge distribution on *P. falciparum* and the erythrocyte membrane makes a compelling model where peptides of either charge, negative or positive, may interfere with parasitic invasion and infectivity (***24***).

In this current study, we demonstrate a strategy of using the parasite’s advantage in making long poly-lysine peptides into their weakness. Under the assumption that poly-basic residues are involved in host cell attachment and invasion, we aimed to utilize poly-lysine peptides for clearing erythrocytic *Plasmodium* with the rationale that treatment of infected cells with poly-basic peptides may block invasion. As such, we use exogenously added poly-cationic peptides to illustrate a specific growth inhibitory effect against *P. falciparum* in human erythrocytes *in vitro*. Our results show that the effects of poly-lysine peptides depend on the length of the peptide, its stereochemistry, and the link between lysine residues in homo-polymers. Our results indicate potent and similar lethal effects of 25-30 residue long α-poly-L and poly-D-lysine peptides on *P. falciparum* with no effect of ε-poly-lysine peptides. Coupling of poly-lysine with non-peptide moieties, as well as the use of poly-basic branching dendrimers, show an increase in the effectivity of poly-lysine peptide-based therapies against *P. falciparum in vitro* with no obvious cytotoxic effects towards human tissue cultures. Using fluorescently-labeled and biotinylated poly-lysine probes for live-cell imaging, we demonstrate that the effects of poly-lysine peptides are associated with specific interactions with the membranes of infected erythrocytes or late-stage erythrocytic parasites. Finally, our pull-down assays from infected erythrocytes on a poly-lysine matrix coupled with mass-spectrometry-based proteomic analyses indicate a set of *P. falciparum* merozoite and erythrocyte membrane and adhesion proteins as the main targets of poly-basic peptides. Our results demonstrate a novel route forward for the potential development of new and powerful peptide-based therapeutics against malaria parasites.

## Results

### Polycationic peptides inhibit *Plasmodium falciparum* growth

Recent attention has been drawn to poly-lysine due to its abilities in drug delivery, antimicrobial activity, biocompatibility, anti-cancer activity, and prion propagation prevention (***15, 25***). Sulfated polyanions, especially heparin **(Fig. 1A)**, have been shown to inhibit the invasion of erythrocytes by merozoites (***22, 26***), with an IC50 for heparin noted as 11 µg/ml (***22***). To test the potential activity of polycationic peptides against *P. falciparum* parasites, we compared the activity of poly-L-lysine (PLL) and poly-L-ornithine with heparin and poly-L-glutamate in a single dose, 72 h *in vitro* treatment experiment **(Fig. 1B)**. We utilized poly-amino peptides in mixtures of 20-50 amino acid residues with molecular weights ranging from 3.5 to 6.5 kD. A control set of parasite cultures were incubated in the absence of any compound or in the presence of PEG-3550 as an inert and non-charged compound. The activity exhibited from PLLs was comparable to that of heparin - more than 50% inhibition of parasite growth at 12.5 µg/ml for both compounds **(Fig.1 B)**. Poly-ornithine demonstrated similar activity as PLL peptides. At the same time, we did not observe parasite growth inhibition in the presence of either poly-L-glutamate or PEG-3550, arguing that the charge and structural elements of poly-cationic peptides or heparin play an important role in this activity **(Fig.1 B)**.

**Figure 1.**
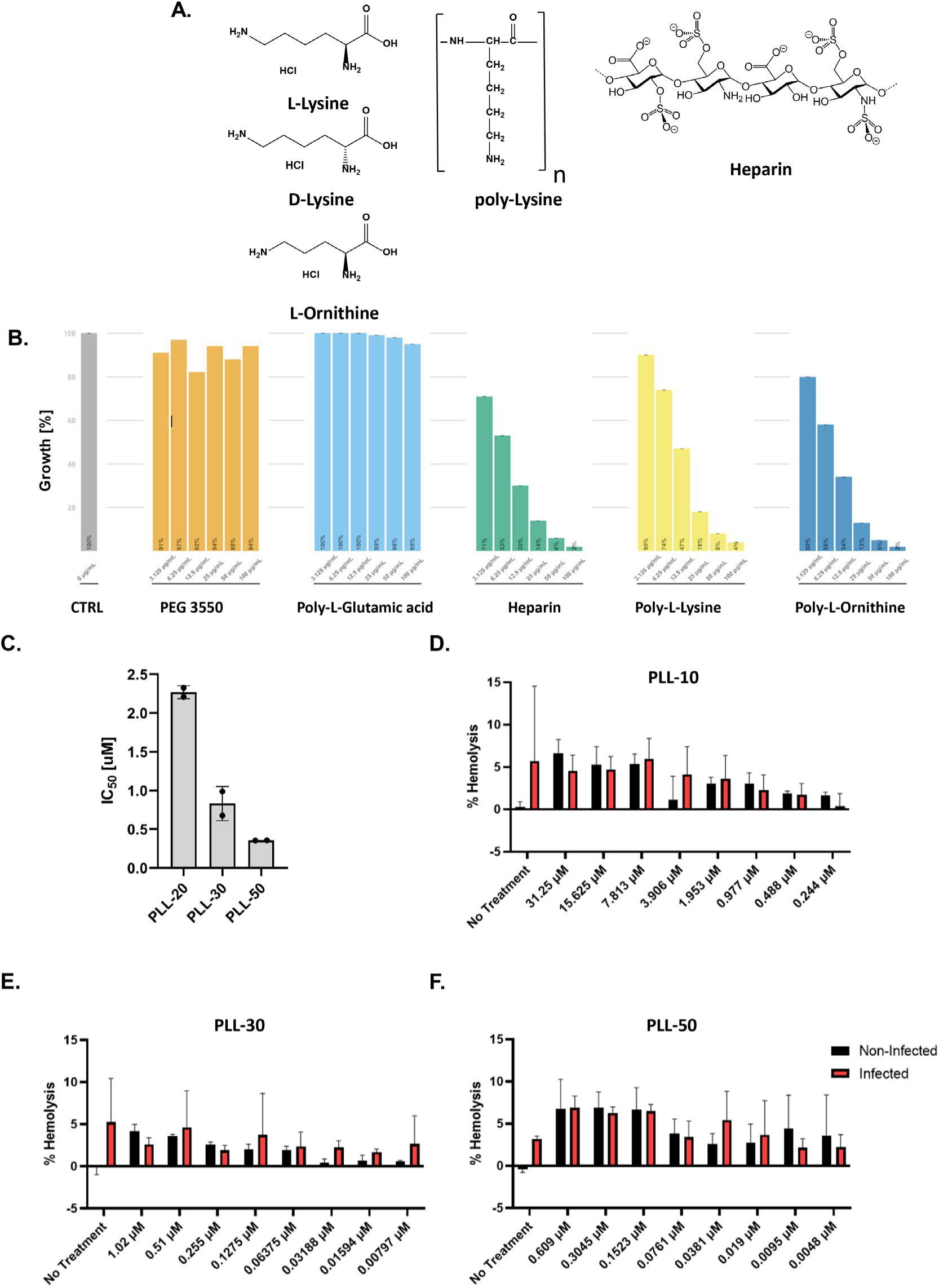
In vitro anti-Plasmodium effect of positively charged peptides and their impact on erythrocyte’s lysis. **A**. Structure of L-Lysine and D-Lysine and an indication of link in poly-Lysine polymers (n - depicts a different number of residues in homopolymers), L-Ornithine, and heparin. **B**. *P. falciparum* Dd2 strain growth assay in the presence of different compounds and amino acid polymers. Data represent endpoint parasitemia after 72-hour treatment with the indicated compound. Results are normalized to non-treated cultures (control) and presented as a percentage of control samples. The average value from three biological replicates is reported with a standard deviation. **C**. IC50 values for different poly-L-lysine homo-polymers (PLL). PLL-20, PLL-30, and PLL-50 treat *P. falciparum* Dd2 strain. Results are normalized to non-treated cultures, and the graph presents two biological replicates done in triplicates with a standard deviation. **D, E, F.** Effects of PLL-10, PLL-30, PLL-50 on lysis of *P. falciparum*-infected and non-infected erythrocytes (black) (three biological replicates, with standard deviation).

Following our experiments on parasite growth inhibition of mixed PLL peptides with different lengths (mixture of peptides with 20-50 lysine residues long; molecular weight 3.5-6.5 kDa **(Fig.1 B)**), we investigated whether the length of poly-L-Lysine peptides plays a role in the efficiency of *P. falciparum* growth inhibition. We utilized PLLs of defined lengths (20, 30, and 50 consecutive lysine residues) and followed the survival of the *P. falciparum* Dd2 strain after 72 hours incubation with the different compounds **(Fig. 1C)**. The IC50 values for each compound were obtained using acridine orange staining **(Fig. 1C)**. The most potent inhibition was achieved in the presence of PLL-50 and calculated IC50s of 0.355 ± 0.0014 µM, PLL-30 followed with IC50 of 0.83 ± 0.221 µM and PLL-20 IC50 of 2.2 ± 0.155 µM **(Fig. 1C)**. The differences in activity indicated that the length of the PLL peptides ostensibly impact might affect *P. falciparum* growth inhibition.

To determine if the impact of PLLs on parasite growth was due to erythrocyte lysis, we conducted a 72-hour incubation of both non-infected and infected human erythrocytes with increasing quantities of PLL-10, PLL-30, and PLL-50 (**Fig. 1D**). We tested hemoglobin levels released from the lysed erythrocytes at the beginning of the experiment and after 72 h for both infected and non-infected cells. The positive control was sampled with total lysis of erythrocytes, and the negative control was sampled with erythrocytes in the culturing medium without any additional compounds. The effect of PLLs on erythrocyte lysis was insignificant, as most erythrocyte cultures exhibited less than 5% lysis (**Fig. 1D**). Moreover, there was no notable difference between infected or non-infected erythrocytes. As such, our results indicate that poly-L-lysine peptides can inhibit the growth of the *P. falciparum* parasites without affecting host cells.

### Poly-lysine compounds are efficient against artemisinin and chloroquine-resistant *P. falciparum* strains

As the resistance to artemisinin and ACT therapy is rising in *P. falciparum* isolates worldwide, we wanted to test the effect of poly–lysine peptides on different strains of *P. falciparum* - specifically the strain that is resistant to artemisinin. We tested the effect of poly-lysine on the artemisinin-susceptible (MRA-1239) and the resistant (MRA-1238) strains (***27***). We utilized peptides with runs of 30 and 50 consecutive lysines (PLL-30 and PLL-50) and the more protease-resistant poly-D-lysine peptide (PDL-30). Both PLL-30 and PLL-50, as well as PDL-30 and PDL-50, had comparable effects on the growth of *P. falciparum* parasites **(Fig. 2)**. Growth of the MRA-1239 strain was inhibited with an IC50 of approximately 11 ng/ml of poly-lysine peptides (**Fig. 2A**). In comparison, the artemisinin-resistant strain (MRA-1238) was inhibited at an IC50 of 8 ng/ml (**Fig. 2B**). We also compared the effects of poly-D-Lysine (PDL-30), poly-L-Lysine (PLL-30 and PLL-50), and artemisinin (ART) on the growth of chloroquine-resistant (Dd2) and non-resistant (NF54) strains of *P. falciparum* **(Fig.2 C** and **D).** The IC50s values for the chloroquine-resistant Dd2 strain were consistently lower than those of the chloroquine-sensitive NF54 strain, with average values of 6 µg/ml and 12 µg/ml, respectively. However, we did not observe a difference between PLL-30 and PDL-30 peptides, as both were equally effective and active against the chloroquine-resistant (Dd2) and chloroquine-susceptible (NF54) strains. Finally, we assayed the toxicity of the PLL-30 peptides utilizing cultured HepG2 cells (**Fig. 2E**). We found that PLL-30 peptides did not exhibit any additional toxic effects compared to control conditions (addition of DMSO) in the range of their observed activity against malaria parasites. As such, our experiments indicated that despite the differences in the chirality of the lysine residues, L versus D, PLL, and PDL peptides had the same growth inhibition effect on different *P. falciparum* strains. Moreover, the inhibitory effect was similar, regardless of parasite resistance to artemisinin or chloroquine, arguing for novel parasite targets by poly-lysine peptides.

**Figure 2.**
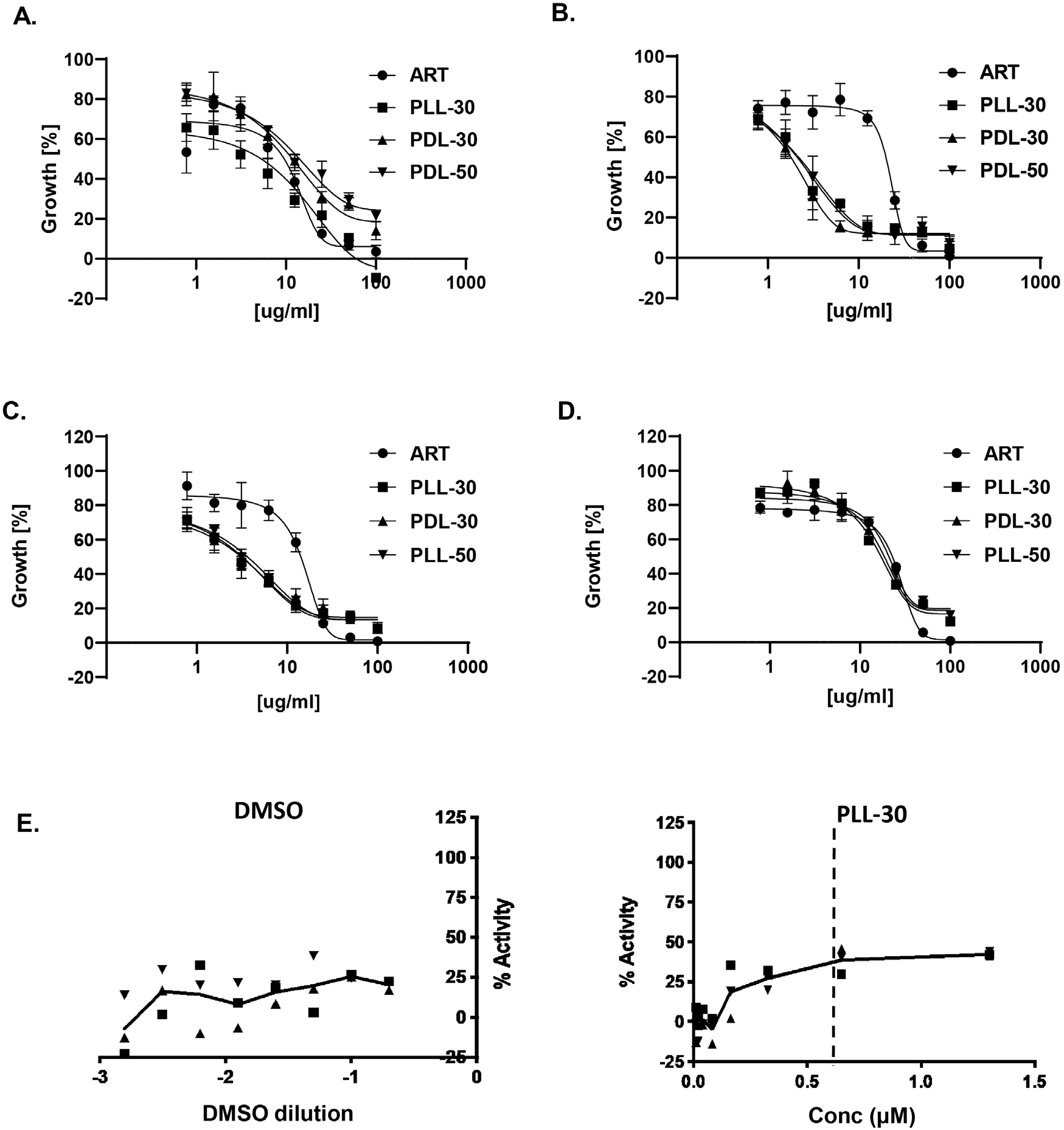
Different forms of poly-lysine have the same activity on artemisinin or chloroquine-resistant and non-resistant P. falciparum strains with low human cell cytotoxicity. **A, B.** Effects of poly-L-Lysine (PLL-30) and poly-D-Lysine (PDL-30 and PDL-50) compared to (●) artemisinin (ART) on the growth of *P. falciparum* artemisinin-resistant and non-resistant strains, respectively (three biological replicates, with standard deviation). **C, D.** Effects of poly-L-Lysine (PLL-30) and poly-D-Lysine (PDL-30 and PDL-50) compared to (●) artemisinin (ART) on the growth of P. falciparum chloroquine-resistant (Dd2) and non-resistant strains (NF54), respectively (three biological replicates, with standard deviation). **E.** Cytotoxicity of PLL-30 within HepG2 cells. Cell viability was presented as a % normalized of untreated control, and dilutions of DMSO were used as a control for dilutions of drugs (three biological repeats).

### Modifications of poly-lysine peptides further inhibit the growth of *P. falciparum*

Previous research has indicated that PEG modification of peptides (PEGylation) can enhance the effectiveness of peptide-based therapies (***16***). PEGylation, the covalently attaching PEG molecules to another molecule (often a drug or protein), can improve the solubility, stability, and half-life of the drug or protein (***28***). To test the effect of PEG modifications on poly-lysine peptides and their ability to inhibit the growth of *P. falciparum* in erythrocytes, we first compared PLL-30 peptides with PEG-PLL50 (A-B connection of PEG (B) to peptide (A)) and PLL-50-PEG-PLL-50 (A-B-A connection) copolymers (**Fig. 3A**). The IC50s for all three compounds were in the sub-micromolar range. The activity of the PEGylated peptides was, on average, three-fold higher with associated IC50s around 250 nM, while the IC50 for PLL-30 was at 800 nM. We also compared *P. falciparum* growth inhibition in the presence of shorter poly-lysine and PEG copolymers (A-B-A; PLL-10-PEG-PLL-10) **(Fig. 3B).** We utilized previously tested PLL-50, PEG-PLL50, PLL-50-PEG-PLL-50 and poly-epsilon-lysine (poly-ε-lysine) peptides. PLL-50 and the, along with longer PEGylated poly-lysine copolymers (PEG-PLL50 and PLL-50-PEG-PLL-50), with all having similar effects on parasite growth inhibition; 50% inhibition at 11µg/ml (IC50s in the range of 80-1030 nM). A shorter PEGylated poly-lysine copolymer (PLL-10-PEG-PLL-10) was less effective, with 50% growth inhibition at 5.5ug/ml (IC50 in the range of 1µM). We did not observe any parasite growth inhibition using the poly-ε-lysine peptides (**Fig. 3B**). As in the case of non-modified PLLs, the HepG2 toxicity assays indicated that PEGylated PLL copolymers have relatively low toxicity in the concentration range where they exhibit potent parasite growth inhibition (<400nM). However, we did notice that higher toxicities were associated with high concentrations of PEGylated PLL copolymers (>1µM) when incubated for 72 hours with HepG2 cells (**Fig. 3C**).

**Figure 3.**
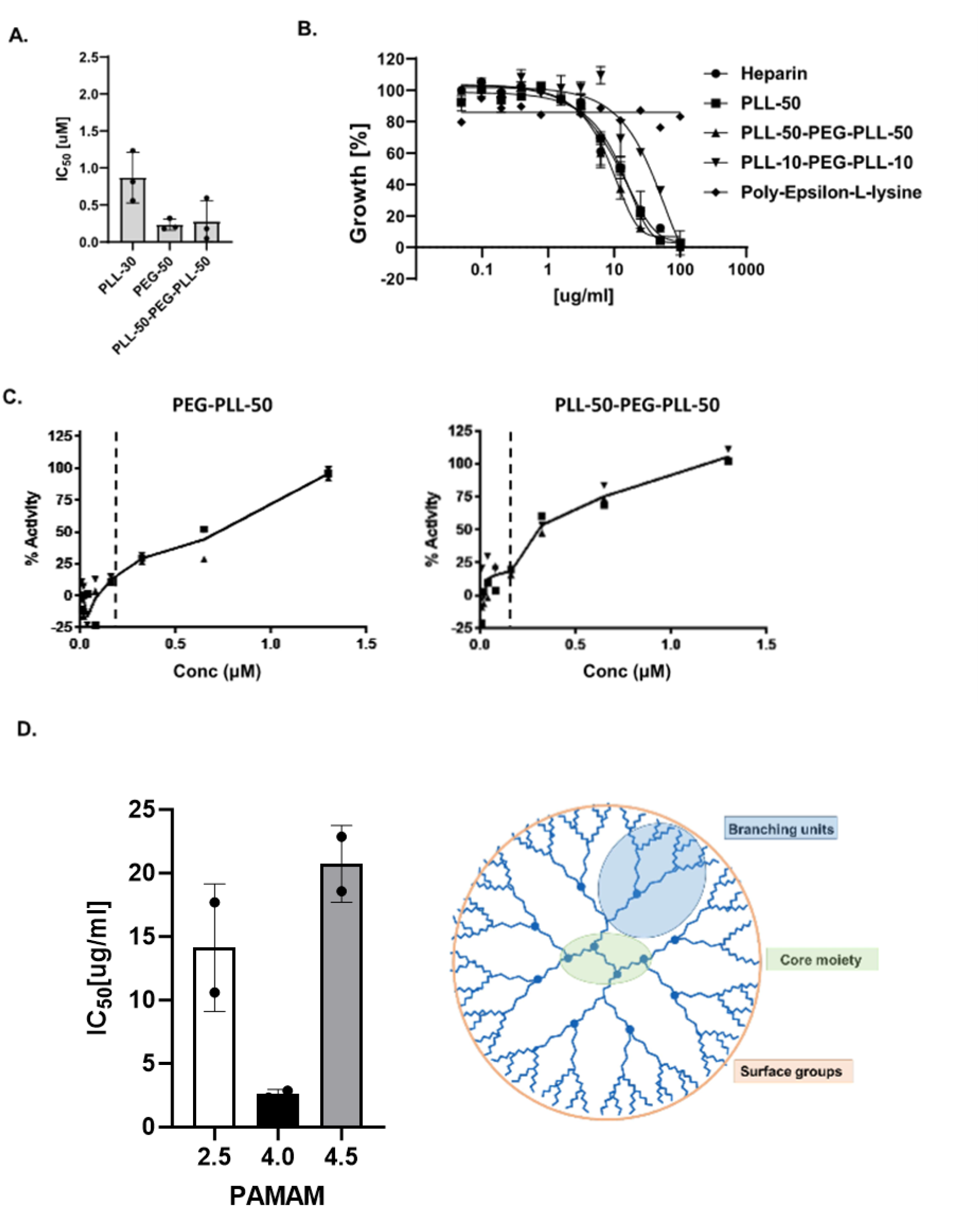
Modifications of poly-Lysine as well as charged polyamidoamine (PAMAM) dendrimers indicate inhibition of *P. falciparum* growth in erythrocytes with low human cell cytotoxicity. **A.** IC50 values for non-modified (PLL30) and PEG-ylated forms of PLL (PEG-PLL-50 and PLL-50-PEG-PLL-50) on *P. falciparum* Dd2 strain growth. **B.** The effect of non-modified (PLL-50), PEGylated (PLL-10-PEG-PLL-10; PLL-50-PEG-PLL-50), and epsilon-linked poly-L-Lysine on *P. falciparum* parasites growth. Heparin was used as a positive control (three biological replicates, with standard deviation) **C.** Cytotoxicity of PLL-50 with PEG modifications on HepG2 cells. Cell viability was presented as a % normalized of an untreated control (three biological repeats are indicated in each graph). The dashed line indicates IC50 values against Plasmodium growth for each tested compound. **D.** IC50 values for polyamidoamine (PAMAM) dendrimers of different generations (2.5, 4.0, and 4.5 generation) used for the treatment of *P. falciparum Dd2* strain (three biological replicates, with standard deviation). Schematics of the PAMAM molecule of generation 4 with characteristics of the molecule are shown (Biorender).

Finally, given that polyamidoamine (PAMAM) dendrimers, as synthetic mimics of branched poly-lysine peptides, have been investigated for a wide range of biomedical applications due to their low cytotoxicity and potential to be further modified, we tested their activity against *P. falciparum* parasites (***29–32***). We investigated three different PAMAM compounds classified as 2.5-, 4-, and 4.5-generation, based on their different architecture, size, shape, and surface functionality (**Fig. 3D**). PAMAM 4.0 generation showed the highest activity in the inhibition of *P. falciparum* growth, with IC50 in the range of 2.5 µg/ml **(Fig. 3D).** PAMAM of 2.5 and 4.5 generation exhibited lower activity in inhibiting *P. falciparum* growth with IC50s in the 13 and 21 µg/ml range, respectively. While this observed difference could be due to the distribution of positive and negative charges between PAMAMs of different generations, our results with both PEGylated or branched modified poly-lysine peptides and co-polymers strongly implicate the role of positively charged polymers in the effective inhibition of *P. falciparum* parasites growth.

### PLL peptides bind to the membrane of parasites and infected erythrocytes

Our previous studies have indicated that *P. falciparum* proteins with poly-lysine repeats are associated with cellular adhesion and are likely to be located at the surface of the parasite (*17*). To assess the mechanism of action and localization of the poly-cationic peptides and co-polymers against the malaria parasite, we designed a fluorescein isothiocyanate (FITC)-labeled 25 lysines residue-long PLL peptide (**Fig. 4A**). In addition to the FITC-labeling we have also engineered HA- and as well as biotin-tagged peptides. We also utilized a previously tested PLL-30 peptide by binding one FITC molecule per each 10-lysine residues. We first tested the activity of the labeled and modified peptides in inhibiting the *P. falciparum* growth for 72 hours. We observed similar trends in parasitic growth inhibition when FITC-HA-PLL-25-BIO or FITC-labelled PLL-30 peptides were incubated with parasites, albeit without complete 100% inhibition (90% inhibition achieved) at the working concentrations **(Fig. 4A)**. Imaging of the parasite cultures in the presence of the FITC-labeled peptides indicated that the poly-lysine peptides were bound to the membrane of infected erythrocytes or parasites (**Fig. 4B-C**, **Supplementary Fig. 1**). Moreover, labeled PLL peptides were bound to specific developmental stages of *P. falciparum* parasites - the surface (**Fig. 4B**) as well as the late schizont stage **(Fig. 4C)**. The labeled peptides did not bind to uninfected erythrocytes or human tissue cultures used as controls (**Fig. 4B**-**C**, **Supplementary Fig.1**) highlighting their specificity towards *P. falciparum* targets or modifications to the human host cell membrane influenced under *P. falciparum* infection.

**Figure 4.**
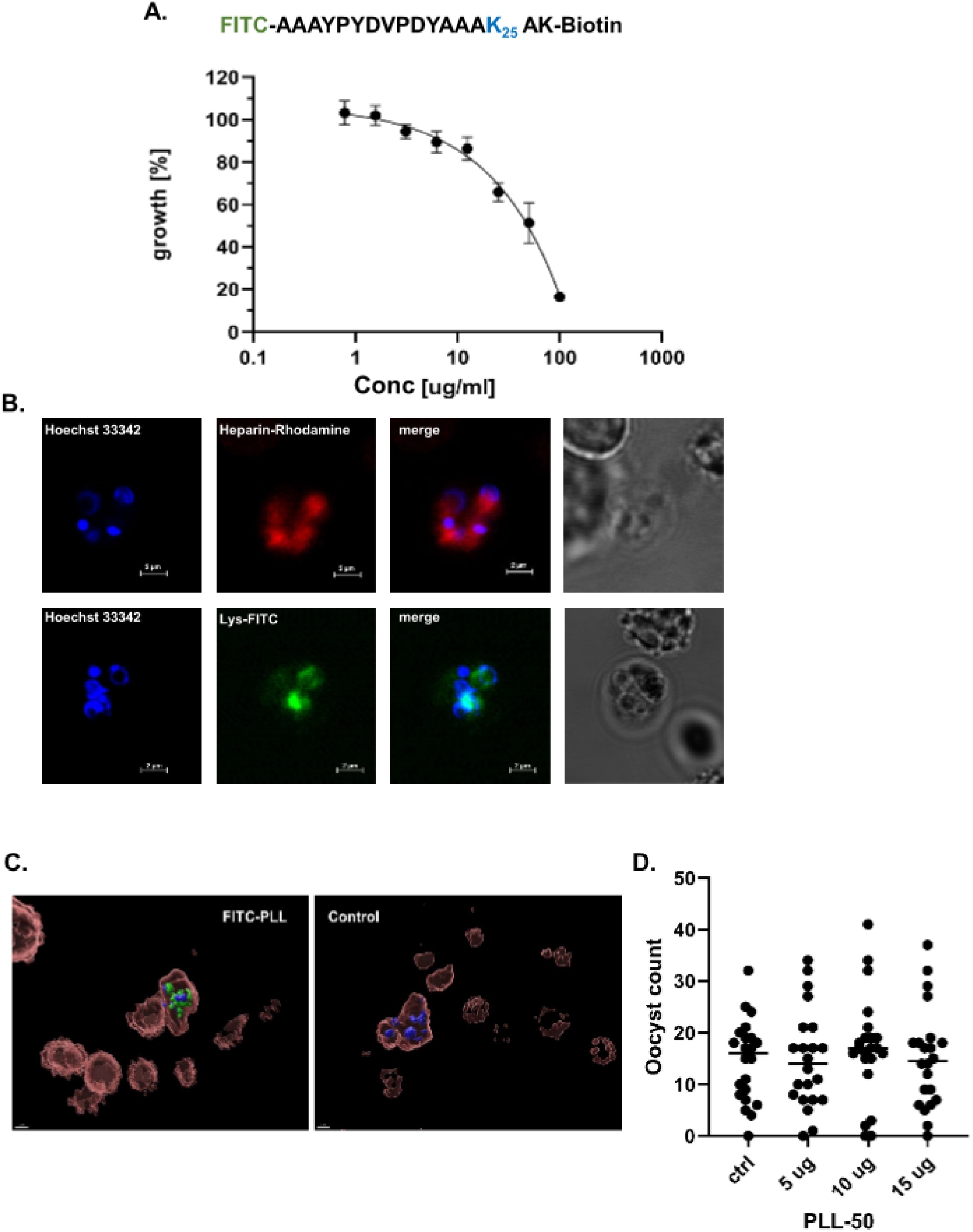
Biotinylated and fluorescence (FITC) labeled poly-lysine peptides indicate the activity of polybasic peptides at membranes of *P. falciparum* in a stage-specific manner. **A.** Inhibition of *P. falciparum Dd2* growth using biotinylated and FITC labeled poly-lysine (FITC-HA-PLL-25-BIO (three biological replicates, with standard deviation, are presented in each experiment). **B.** Rhodamine-labeled heparin and FITC-PLL binding to Plasmodium merozoites. The FITC-PLL-Biotin indicates membrane binding on merozoites. Hoechst 33342 is used for nuclear staining. The bar indicates 2 um length**. C.** Airyscan confocal fluorescence imaging microscopy of FITC-PLL-Biotin treated (FITC-PLL) and non-treated *P. falciparum* Dd2 erythrocytes at late stages of parasite development. Hoechst stain was used to visualize *P. falciparum* nuclei. Bar represents the size of 2um. **D.** Effect of PLL-50 on oocyst survival. Counts of viable oocysts between control (ctrl) and treated samples (5,10 and 15 ug per ml of PLL-50) do not show any significant change.

Since the *Plasmodium* life cycle involves parasite development in mosquitos, we also tested the influence of PLL on parasite growth and development in mosquito guts. We did not see any PLL effect on the oocyst parasite growth stage in mosquito hosts **(Fig. 4D)**. However, imaging of FITC-HA-PLL-25-BIO peptides incubated with *P. falciparum* parasites isolated from mosquito salivary glands indicated that PLL peptides could interact with sporozoite membranes **(Supplementary Fig. 2).** The binding of FITC-HA-PLL-25-BIO peptide to sporozoites had a similar distribution as previously seen in imaging from parasite merozoites and schizonts. Our results using FITC-labeled PLL peptides indicate that PLL peptides can bind specifically to the membrane of *P. falciparum* parasites or infected cells across multiple parasite developmental stages.

### Poly-lysine peptides bind to a subset of *P. falciparum* membrane proteins, potentially inhibiting erythrocyte invasion

As poly-lysine peptides could affect the function of many different proteins, we worked to assess the potential targets of poly-lysine and poly-cationic peptides in *P. falciparum* and parasite-modified human host cells. We incubated the poly-L-lysine agarose, PLL chains with 30-70 residues, with cell lysates from erythrocytes either infected or uninfected with the *P. falciparum Dd2* strain using non-infected erythrocyte lysate as control (**Fig. 4A**). Infected erythrocyte cultures were previously developmentally synchronized to increase the number of parasite late schizonts and merozoites from the parasites. Using mass-spectrometry **(Fig. 4B)**, we identified 1295 proteins from either human or parasitic cells (**Supplementary Table 1**). Of these, 950 proteins were identified with more than two peptides in spectral counts (**Fig. 4C**, **Supplementary Table 2**). Six hundred forty-five proteins showed an increase in infected erythrocytes compared to non-infected control (**Supplementary Table 2**). Most of these proteins were *P. falciparum* proteins associated with various cellular processes, with 423 proteins enriched over 1.5-fold (a total of 473 *Plasmodium* proteins, **Supplementary Table 2**). Since our previous experiments and incubation of the parasite cultures with FITC-labeled PLLs indicated membrane association of the charged peptides, we focused our analyses on the *P. falciparum* membrane and surface proteins. We were able to identify, based on gene annotation, 75 *P. falciparum* exported and membrane-associated proteins, from which 57 proteins had significant P-values and enrichment over 3-fold (**Fig. 4C** and **Supplementary Table 2**). These *P. falciparum* proteins were associated with merozoite invasion, host cell surface expression and interaction, and vacuole membranes and extracellular granules. In addition to *Plasmodium* proteins, we identified 172 human host proteins, of which 134 were enriched over 1.5-fold in *Plasmodium*-infected cells. Finally, we identified 46 host membrane-associated proteins, of which 25 proteins had significant P-values and enrichment over 3-fold within host cells **(Supplementary Table 2**). Considering our data using FITC labeled PLLs and LC-MS experiments on the PLL matrix, we can indicate multiple surfaces and membrane-associated proteins of *P. falciparum* as potential targets of poly-lysine and other poly-cationic polymers.

## Discussion

Anti-microbial peptides (AMPs) have been used as potential therapeutic compounds for multiple single-cell microbes and viruses (***11***). AMPs are usually composed of 15-60 amino acids. They can be naturally expressed in multicellular organisms or synthetically made and typically possess a positive charge under physiological pH conditions (***33***). As such, the activity of AMPs are usually due to their ability to interact with the negatively charged components of bacterial cell membranes (***11, 12, 14***). This interaction can lead to cell membrane disruption, resulting in cell death. The complex structures of AMPs allow them to interact with microbial membranes more efficiently or with higher specificity, and they often contain additional functional elements that can enhance their antimicrobial activity against specific microbes (***9, 13, 34***). One such group of naturally- and synthetically-made AMPs represent poly-lysine peptides. Regardless of their link position, α or ε, poly-lysine peptides are potential antimicrobial agents against multiple bacterial species, yeast, or fungi (***35, 36***). Interestingly *P. falciparum*, and *Plasmodium spp.* in general, contain numerous poly-L-lysine stretches in proteins that are involved in the parasites’ life cycle functions, mainly focused on cell adhesion and pathogenicity (***17***). Previous work exploited heparin, a negatively charged glycosaminoglycan, to interfere with *P. falciparum* invasion (***22, 23***). Furthermore, charge distribution and polarization of *P. falciparum* membrane during invasion make a compelling case to test peptides of different charges against parasites (***24***). Given that the range of AMPs has shown potential activity against malaria parasites and their vector mosquitos, here explored the possibility of targeting *P. falciparum* parasites with different poly-lysine and poly-cationic polymers.

Previous work from multiple labs has focused on inhibiting *P. falciparum* growth or development using naturally occurring or complexed synthetic peptides (***10***) with a mix of charged and hydrophobic residues. Our results indicate a potent anti-malarial activity of compounds based on simple poly-lysine or poly-cationic polymers with additional non-amino acid modifications (**Fig. 1-4**). These compounds show sub-micromolar potency against *P. falciparum* asexual blood stage and complete inhibition of further growth in a single dose treatment of *in vitro* parasite cultures in human erythrocytes. The activity of poly-lysine peptides is not dependent on the chirality of the lysine residues (L or D). Still, the activity of poly-lysine peptides is associated with peptide length and residue linkage (**Fig. 1-2**). Our results indicate that peptides with a length of 25-50 lysine residues and an α-link between the lysine residues exhibited robust anti-malarial activity. These synthetic poly-lysine peptides, PLLs, and PDLs inhibit the growth of multiple *P. falciparum* strains, including one with known artemisinin resistance (**Fig. 2**), the compound in the current therapy of choice to treat malaria infection. Importantly, we found no cytotoxic effects of poly-lysine peptides on cultured HepG2 cells. Nor did we see the increased hemolytic activity of PLLs at concentrations that cause efficient inhibition of parasite growth. Cytotoxicity and hemolytic activity of antimicrobial peptides has previously been described as a reason for the limited use of such compounds in treating various infections and diseases (***37, 38***).

Our results also indicate that the naturally produced antimicrobial peptide and common food preservative poly-ε-lysine did not exhibit any anti-malarial activity (**Fig. 3**). This is in contrast to studies on Gram-positive and -negative bacteria, where ε-poly-L-lysine shower far greater bactericidal activity than α-PLL (***39–42***). This discrepancy is likely due to the different structures and distinct modes of action of these poly-lysine polymers, as previously observed in evaluating multiple poly-lysine polymers (***43***). Poly-L-lysine’s activity might be relatively low compared to other more complex or modified antimicrobial peptides (***44–47***). Amphiphilic compounds based on partially PEGylated poly-lysine peptides may offer several advantages, such as reduced cytotoxicity, the induction and stabilization of self-assembled nanoparticles, and extended *in vivo* circulation time. These nanoparticles demonstrated comprehensive antimicrobial activity against clinically relevant gram-positive and gram-negative bacteria while maintaining minimal hemolytic activity (***48***). Our results using PEGylated poly-lysine co-polymers illustrate the growth inhibition of *P. falciparum* blood stages at lower IC50 concentrations. PEGylated PLL50 or copolymer of PEG with two PLL50 peptides showed IC50s 4 to 8 times lower than non-PEGylated PLLs (**Fig. 1-3**).

In addition to PEGylation, further efforts were made to mimic poly-L-lysine polymers or combine them to enhance their antimicrobial activity and stability. These included combinations of charged non-amino acid moieties or the use of branched lysine or non-lysine polymers such as dendrimers (***15, 44***). A series of such compounds are already used in multiple biomedical applications, including drug delivery and, more specifically, as vehicles for gene therapy (*29, 45, 49, 50*). Our experiments tested three generations of poly-amidoamine, or PAMAM, dendrimers with different mixtures of amine and carboxyl-terminal groups. We utilized PAMAMs of 2.5, 4, and 4.5 generations, where a half-generation number indicates that only one type of reaction has been carried out: amination or methylation. Our results indicate that PAMAM dendrimers of generation 4, where four complete generation cycles are carried out, resulting in a larger, more complex molecule with more terminal amino groups present on the surface, demonstrate the best growth inhibition against P. falciparum parasites (**Fig. 3**). While PAMAM 2.5 and 4.5 showed reduced activity, likely due to the reduction in the number of charged surface groups, the potential for further. The variability in modifying these molecules makes them attractive candidates for further investigation as anti-malarial drugs.

Previous studies targeting malaria parasites with antimicrobial peptides have indicated activity for some of the tested peptides, mainly in micromolar ranges (***10***). However, these studies did not define specific target molecules or the mode of action of tested peptides but rather reported developmental stages influenced by active peptides (***10***). Given that most antimicrobial peptides act at the cell wall or the membranes of the targeted microbes, we were interested in how poly-lysine and poly-cationic peptides target and act on *P. falciparum.* We utilized fluorescently-labeled poly-Lysine (FITC-PLLs) as well as fluorescently-labeled and biotinylated HA-tagged PLLs (FITC-HA-PLL-25-BIO) peptides to target blood stages of *P. falciparum* (**Fig. 4 and Supplementary Fig 1**). Incubation of these labeled and modified PLLs exhibited similar and effective growth inhibition against the tested *P. falciparum* Dd2 strain. Moreover, by imaging parasite cultures in the presence of FITC-labelled PLLs, we noted that the binding of peptides was specific for the parasites in the merozoite and late schizonts stages of the asexual cycle. The FITC-PLLs did not affect the development of the parasites in the mosquito stage as tested oocytes did not show any reduction in number (**Fig. 4D**). However, we did observe that fluorescently labeled peptides bind to the sporozoite stage parasites isolated from mosquito salivary glands (**Supplementary Figure 2**). While FITC-PLLs had a similar pattern of membrane binding, as seen in merozoites, the effect of PLLs on sporozoites and later stages of parasite growth needs to be evaluated by further study.

Previous work using either PLL or PDL with or without FITC labelling indicated that positively charged peptides neutralized the negative charge of the cell surface in different cells (***49, 51***). We noted that PLLs might have a similar effect on *Plasmodium* membranes as FITC-labeled peptides exhibited similar binding patterns to parasite membranes at different stages. However, we wanted to further identify potential protein targets of PLLs on the surface of infected erythrocytes or the *Plasmodium* membrane. Incubating cell lysates of erythrocytes infected with *P. falciparum* on a poly-L-lysine matrix enabled us to identify a subset of highly enriched parasite proteins (**Supplementary Table 2**). While both cytoplasmic and membrane proteins were enriched as identified via LC-MS analysis, due to the observed membrane binding of FITC-PLLs, we focused on secreted, adherent, and membrane-inserted proteins of *Plasmodium* in the obtained data (**Supplementary Table 2**). Amongst these proteins, the most significant and potential targets of PLLs were Clag9 (PF3D7_0935800), expressed on the merozoite surface and involved in merozoite invasion of human erythrocytes, PIHSTb, c (PF3D7_0401800, PF3D7_0801000 respectively) which are; possible vaccine candidate, GARP (PF3D7_0113000; involved in programmed cell death), EMP1-trafficking protein (PTP4; PF3D7_0730900; required in virulence and rigidity), and PIESP2 (PF3D7_0501200) a; considered as a possible vaccine candidate (***52–56***). Furthermore, additional identification of MDR1 and Kelch13 (***57, 58***) in LC-MS experiments as potential targets of PLLs indicates further possibility to address the resistance to the ACT therapies, as mutations in these proteins give rise to artemisinin resistance. Finally, some of the shorter poly-lysine peptides may penetrate the parasite cells and interact with MDR1 and Kelch13 (***57, 58***), as seen from our LC-MS experiments. Such interactions and possible effects on these targets could address the resistance towards ACT therapies as mutations in these proteins give rise to artemisinin resistance.

Based on previous studies with heparin (***22, 23***), our PLL pull-down, enrichment of the subset of proteins involved in erythrocyte invasion, those exposed on infected erythrocyte membrane surfaces, as well as late schizont and merozoites stage utilizing fluorescently labeled PLL, we propose a mechanism of action for poly-lysine and poly-cationic peptides against *P. falciparum* (**Fig. 5D**). The positively-charged peptides appear to bind to the predominantly negatively-charged body of the merozoite, similar to that of the binding of heparin to the apical region of merozoites (***22–24***). Such a binding mechanism may inhibit or delay the invasion of human erythrocytes by the parasite, which consequentially leads to parasite death. As such, poly-cationic peptides could impede the progression of the infection and result in potential clearance of parasites from the host. In addition to these effects of PLLs and poly-cationic peptides on parasites and infected erythrocytes, PLLs may also act on host proteins presented on uninfected cells that may serve as receptors for lysine-rich *Plasmodium* proteins. Imaging with FITC-labelled PLLs did not show specific labeling of un-infected erythrocytes or other tested human cell cultures. However pull-downs on the PLL matrix indicated enrichment of specific human membrane proteins that were previously shown to be invasion receptors for *Plasmodium.* These additional effects on host membranes could also contribute to the proposed mechanism of PLLs targeting the merozoite stage and inhibiting erythrocyte invasion.

**Figure 5.**
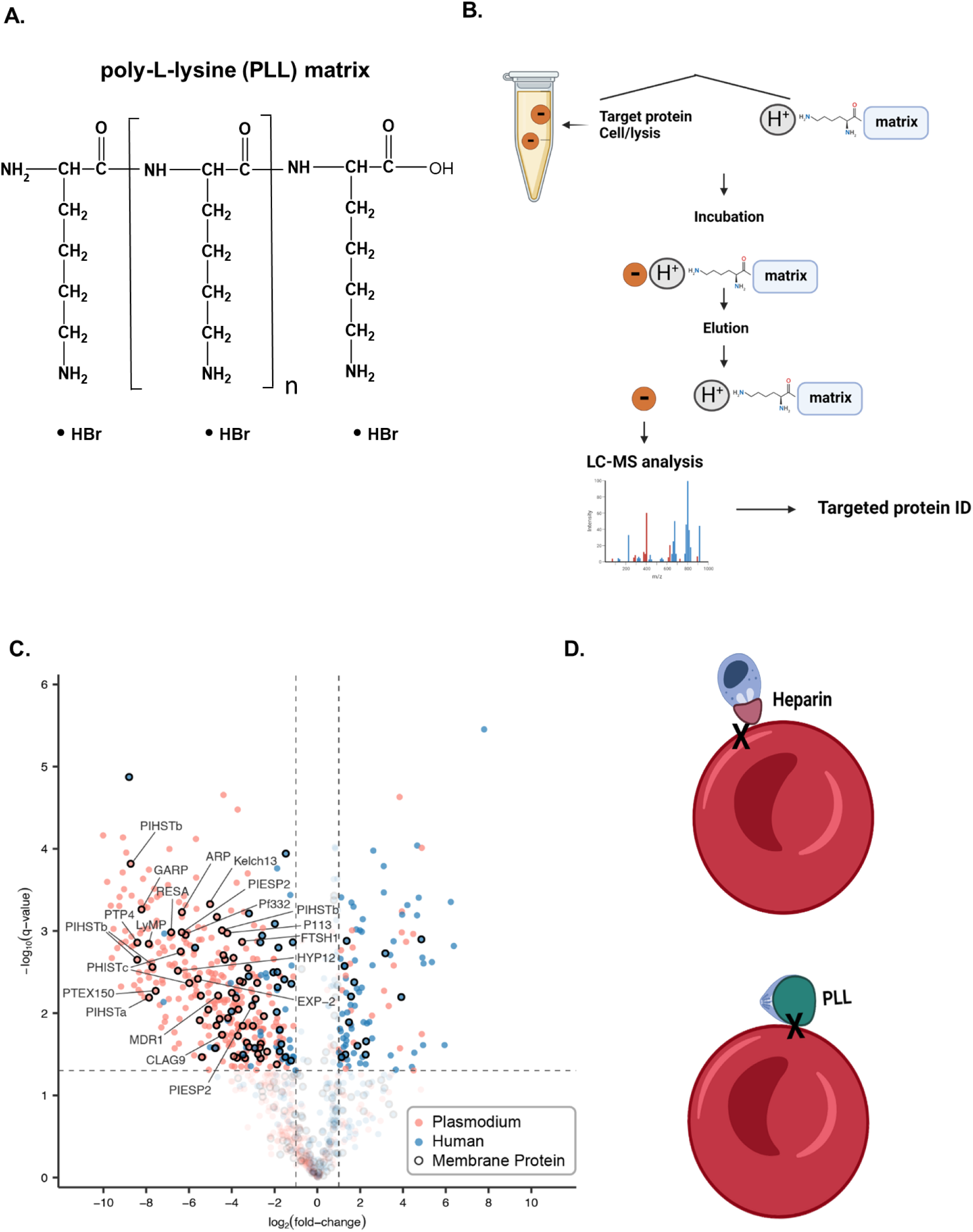
Identification of poly-L-lysine target(s) and proposed mechanism of *P. falciparum* growth inhibition by polybasic peptides. **A**. Poly-L-lysine (PLL) matrix used to identify the target proteins. **B.** Erythrocytes infected with P. falciparum were lysed and incubated with a PLL matrix in tri-biological replicates. Non-infected erythrocytes were used as controls. The proteins bound to the matrix were eluted, and LC-MS analysis was performed to indicate potential targets of poly-l-lysine peptides. **C.** Volcano plot of PLL matrix enriched *P. falciparum* (red) and erythrocytes (blue) proteins from infected and non-infected human erythrocytes. Some of potential targets of PLL peptides at*P. falciparum* cells and plasma membrane proteins are indicated (**Supplementary tables 1 and 2**). Membrane associated proteins from both *P. falciparum* and human erythrocytes are labeled with closed circles. **D.** Proposed mechanism of action for PLL peptides on the invasion of erythrocytes by *P. falciparum*. Positively charged PLL peptides bind to the merozoite body, which is mainly negatively charged compared to the mechanism of heparin binding to the apical part of merozoites. Both compounds act as potential inhibitors of parasite invasion of human erythrocytes.

In summary, we characterized the antimalarial properties of poly-lysine, other poly-cationic peptides, and polymers in this study. We found that a single dose of positively charged peptides can inhibit the growth of *P. falciparum* parasites in human erythrocytes *in vitro*. The poly-cationic peptides and copolymers targeted proteins associated with *P. falciparum* and infected erythrocyte membranes. We hypothesize that the multivalent interaction of charged peptides and related compounds with multiple membrane proteins of *P. falciparum* leads to a reduced number of newly infected erythrocytes and potential clearance of parasites in one or two cycles of asexual replication by preventing new host cell invasion. Our study reveals a potential new direction in creating peptide or peptide mimetic drugs that are potently active against malaria parasite infection.

## Acknowledgments

We are thankful to Goldberg lab at Washington University in St. Louis (Drs. Daniel Goldberg and Eva Istvan) for the help in using the equipment in their lab and for their support. We thank Dr. Peter Agre at Johns Hopkins Malaria Research Institute for his support of this project. Toxicity assays and IC50s for PLL compounds were executed by NYU Langone’s Anti-infectives Screening Core and Rodrigues lab.

## Author contributions

Conceptualization: SD, SPD

Methodology: SD, SPD

Investigation: RS, JE, KF, JKK, AT, MJ, DG, SPD, SD

Visualization: PB, JAJF, AT

Funding acquisition: SD, SPD

Project administration: SD, SPD

Supervision: SD, SPD

Writing – original draft: SPD, SD

Writing – review & editing: RS, JE, KF, PB, JAJF, MJ, AT, DG, SD, SPD

## Data and materials availability

All data, code, and materials used in the analysis are available to any researcher for reproducing or extending the analysis upon contacting corresponding authors.

## Materials and Methods

### Parasite culturing

*P. falciparum* strains (Dd2, Nf54, MRA-1238, MRA-1239) were cultured in deidentified human erythrocytes, maintaining a 2% and 5% hematocrit. The culture medium consisted of RPMI 1640, enhanced with 5 g/l of Albumax II (Thermo Fisher Scientific, 11021037), 0.12 mM hypoxanthine (prepared by adding 1.2 ml of 0.1M hypoxanthine to 1 M NaOH), and 10 μg/mL of gentamicin (***59***). Artesunate (Acros) was used for artemisinin treatment of *P. falciparum* cells. All parasite cultures were maintained statically, with atmospheric conditions provided by a candle jar.

### Parasite sorbitol synchronization

When synchronization was needed, 5% sorbitol (S6021) (by weight/volume) was employed to get the resulting *P. falciparum* culture (primarily ring stage, 5% hematocrit, >3% parasitemia). The resulting parasite culture had to be at least 60% synchronized. If synchronization was below 60%, it was repeated in 6h and 48h to achieve optimal synchronization (***60***).

### Drug Assay

A stock solution for each drug was prepared by diluting 1mL of a 10mg/mL drug solution in sterile water, followed by a 10-minute sonication. For this study we used:

Alamanda Polymers Products:

PLL-20 (000-DKB030-103PLKB20)
PLL-30 (000-KB030-101)
PDL-30 (000-KB050-106PDKB30)
PLL-50 (000-KB050-106)
10-PEG-10 (000-KB020-102PLKC10-b-PEG5K-b-PLKC10)
50-PEG-50 (050-2KC010-101PLKC50-b-PEG1K-b-PLKC50)
010-2KC050-101

Sigma Products:

POLY-L-LYSINE (4000-15000): P-6516
POLY-D-LYSINE (1000-5000): P-0296
POLY-L-ARGININE: P-4663
POLY-D-LYSINE (4000-15000): P-6403
Poly-L-Ornithine hydrobromide: P-3538-50 MG
Poly-L-arginine hydrochloride (10mg): P4663-10MG
Poly-D-glutamic acid sodium salt (1g): P4033
Artesunate (artemisinin) from Acros
Heparin from Creative PEGWorks

Genscript Products:

Rhodamine-AAAYPYDVPDYAAAK_25_: 9.2mg, costume synthesis LOT: U527JGJ060-3/PE6021 FiTC-AAAYPYDVPDYAAAK_25_: 9 MG, costume synthesis LOT: U527JGJ060-1/PE6019

A diluted solution was obtained by combining 392 µL of the designated medium with 8 µL of this stock, which was then chilled on ice. Parasite culture, set at a total volume of 10 mL, was maintained at 4% hematocrit and 1% parasitemia, which was achieved by adjusting a 400 µL sample of 1% parasitemic blood with the necessary medium. For plating, each well of the plate received 100 µL of DSM1 medium using a multichannel pipet. The first row was treated with 100 µL of the diluted drug solution, with distinct drugs in separate columns, and some columns were kept drug-free for control. Serial dilutions were created by transferring 100 µL from one row to the next and discarding 100 µL from the final row to maintain a consistent volume. Each well was then seeded with 100 µL of erythrocytes and incubated at 37°C. Additional erythrocytes were used for smear preparation to examine parasitemia initially and post-incubation if required. Control measures included determining parasitemia at the start (0-hour mark) and in the drug-free wells after 72 hours. Parasitemia analysis after the 72-hour incubation was conducted using flow cytometry.

### Hemolysis Assay

The hemolysis assay was carried out by the previously described protocol (*61*). In the hemolysis quantification protocol, the supernatant was transferred from each well to transparent, flat-bottomed 96-well plates. Triplicate wells, using 4% hematocrit RBCs in water, ensured maximal hemolysis, with Malaria Complete Medium (MCM) serving as the blank control. Absorbance at 574 nm was recorded using the specified instrument. Data was normalized by subtracting mean blank absorbance and further adjusted against the average of the maximum hemolysis controls to derive relative hemolysis percentages. Mean absorbance values were determined for each experimental set from triplicate measurements.

### Cell Labeling

Cells with approximately 5% parasitemia were collected by transferring 100µL into a 1mL microcentrifuge tube, followed by centrifugation at 500g for 5 minutes at room temperature. The pelleted cells were subsequently washed twice with phosphate-free MCM. A staining solution was prepared by diluting 5µL of Hoechst dye in 10 mL of the same phosphate-free MCM. The cell pellet was then resuspended in 1mL of this dye solution and incubated in the dark for 5-10 minutes. After staining, the cells underwent two washes with phosphate-free MCM. For further treatments, a mixture of 10µL of the selected reagent (FITC/Poly-lysine, short-chain poly-lysine, or glutamate) and 990µL phosphate-free MCM was added to the cells and incubated for an additional 5-10 minutes. The treated cells were washed twice with 1mL MCM and resuspended in a final volume of 100µL MCM. CellMask orange plasma membrane stain (Invitrogen) was used for staining of erythrocyte membranes using a concentration of 5mg/mL as 1000x stock solution. Human tissue cultures were incubated with 10µL of the selected reagent (FITC/Poly-lysine, short-chain poly-lysine, or glutamate) and 990µL of opti-MEM for 30 min before imaging.

### Imaging and visualization

Imaging and image analysis were performed at the Washington University Center for Cellular Imaging. Cells were visualized on an upright Nikon A1RHD25 confocal platform (Nikon Instruments, Melville, NY) using a CFI SR HP Plan Apochromat Lambda S 100XC silicone immersion objective. Z-series were collected and reconstructed using Imaris 10.0 (Oxford Instruments, Abingdon, United Kingdom). Cell count analyses and visualizations were performed using the isosurface generation function in Imaris.

Sporozoites were freshly dissected at day 14 post-infection as described earlier (32716382). Live sporozoites were incubated with 1:500 dilution of 100mg/ml FITC labeled Poly-L-Lysine or Rhodamine labeled Poly-L-Lysine for 30 min at room temperature, followed by three washes in PBS. For nuclear staining, 1:1000 dilution of 1mg/ml DAPI (Roche Diagnostics) was included during Poly-L-Lysine labeling. Sporozoites were then viewed with a Zeiss AxioImager M2 fluorescence microscope equipped with an oil-immersion Zeiss plan Apo 100×/NA 1.4 objective or a Zeiss EC plan Neo 40×/NA 0.75 objective and a Hamamatsu ORCA-R2 camera. Optical z-sections with 0.2 µm spacing were acquired using Volocity software (Quorum Technologies, Puslinch, ON, Canada). For siRNA samples, only a single z-plane was acquired. Images were deconvolved with an iterative restoration algorithm using calculated point-spread functions, a confidence limit of 100%, and an iteration limit of 30–35 using Volocity software. Images were cropped and adjusted for brightness and contrast using Volocity software.

### *P. falciparum* NF54 culture and mosquito infection

Mosquito infection with *P. falciparum* NF54 was performed as previously described (32716382). Asexual cultures were maintained *in vitro* in O^+^ erythrocytes at 4% hematocrit in RPMI 1640 (Corning) supplemented with 74 μM hypoxanthine (Sigma), 0.21% (wt/vol) sodium bicarbonate (Sigma), and 10% (vol/vol) heat-inactivated human serum. Cultures were maintained at 37°C in a candle jar made from glass desiccators. Gametocyte cultures were initiated at 0.5% parasitemia and 4% hematocrit. The medium was changed daily for up to 15 to 18 days without adding fresh blood to promote gametocytogenesis. Adult *Anopheles stephensi* mosquitoes (3 to 7 days after emergence) were allowed to feed through a glass membrane feeder for up to 30 min on gametocyte cultures at 40% hematocrit containing fresh O^+^ human serum and O^+^ erythrocytes. Infected mosquitoes were maintained for up to 19 days at 25°C with 80% humidity and provided a 10% (wt/vol) sucrose solution.

### Immuno-precipitation on poly-L-lysine agarose

1 ml parasite pellet was resuspended in 5 mL PBS three times. The pellets were mixed with 2x volume of lysis buffer (150 mM NaCl, 50 mM Tris pH 7.5, 1% IGPAL-CA-630, 5% glycerol, a protease inhibitor (PMSF 1 mM), the samples were left on ice for 15 min. Add 100ul of poly-lysine agarose mixture (Sigma P6893-5 ml, resuspended in 400 ul lysis buffer). The mixture was incubated at 4C for 2 hrs on the rocker. After 2 hrs, the samples were washed with 1 ml wash buffer three times (150 mM NaCl, 50 mM Tris pH 7.5, 5%glycerol. After the washing treatment, the samples were spun down, and the PLL agarose matrix was flesh-frozen and sent for mass spec analysis. The negative control was erythrocyte lysate treated the same as infected erythrocytes.

