## Supplementary figures for "Poly-basic peptides and polymers as new drug candidate against *Plasmodium falciparum*"

### Supplementary Figure 1

A.

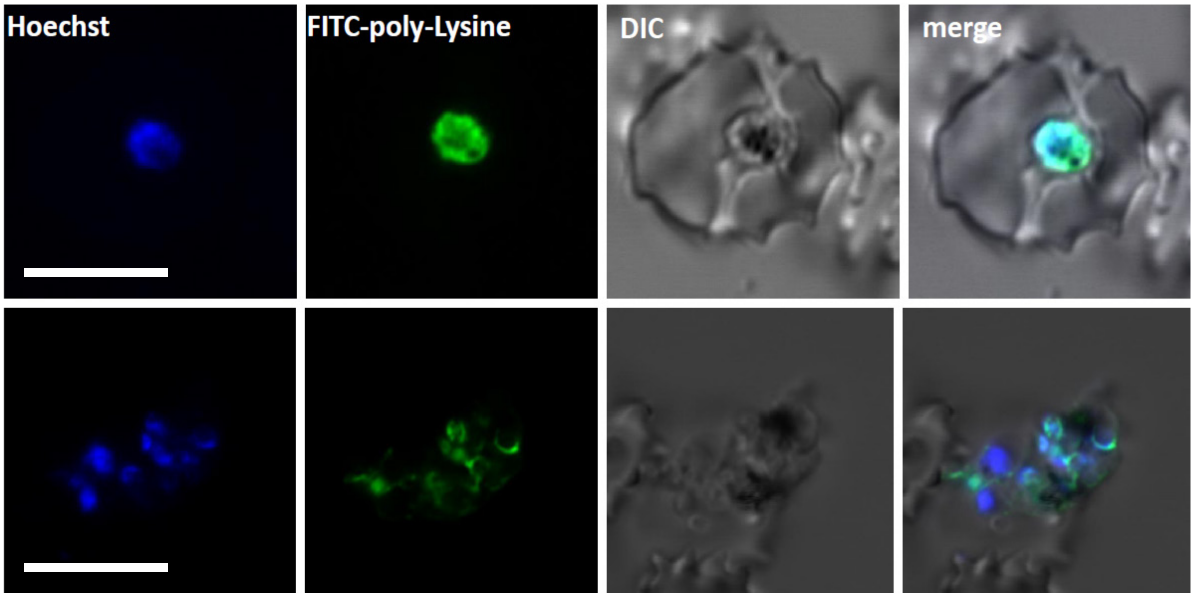


B.

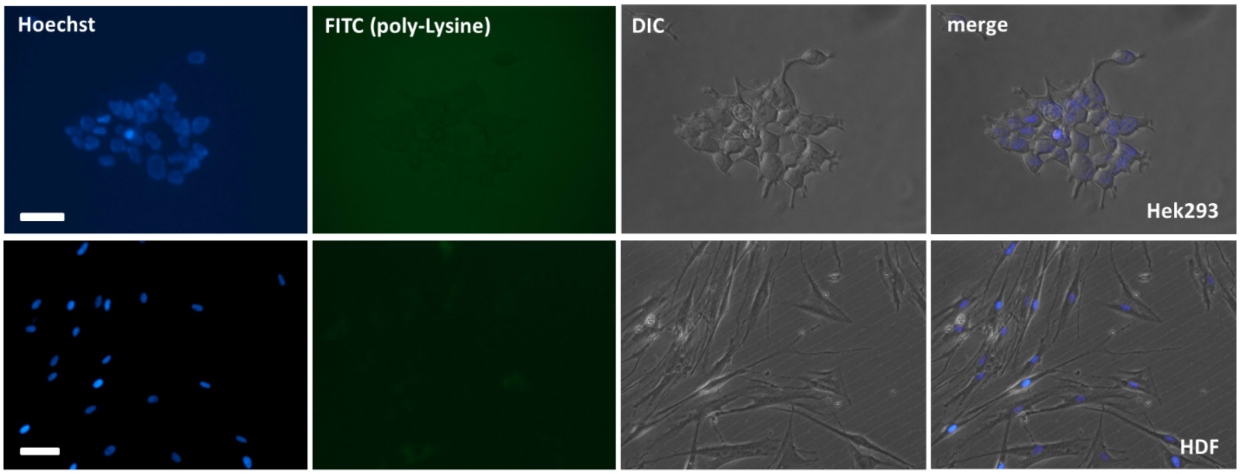


**Incubation of FITC-poly-L-Lysine with infected erythrocytes (A) or human tissue cultures (B). A.** Microscopy images of schizonts and merozoits of *P. falciparum* in infected erythrocytes labelled with FITC-labelled 30 residue long poly-L-lysine. *P. falciparum* membranes are labeled immediately after incubation (1-3 minutes). Bar represents 5µM. **B.** Microscopy images of human tissue cultures, Hek293 and human dermal fibroblasts (HDFs), in the presence of FITC-labelled FITC-labelled 30 residue long poly-L-lysine. Poly-L-lysine incubation of 30min (or longer) does not indicate any visible binding to membranes of human cells. Bars represent 50µM (Hek293) and 30µM (HDF), respectively. Images represent microscopy using different channels (Hoechst, FITC, DIC (optical) and merge, using EVOS imaging system.

### Supplementary Figure 2

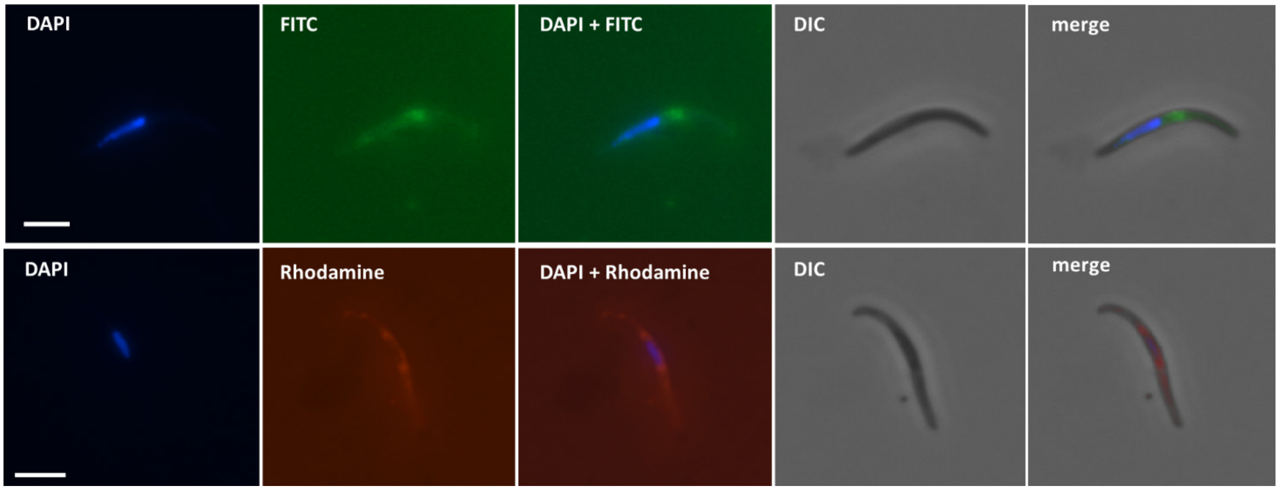


**Incubation of mosquito isolated *P. falciparum* sporozoites with FITC- or Rhodamine-Poly-L-Lysine.** Microscopy images of mosquito isolated sporozoite of *P. falciparum* incubated with FITC- or Rhodamine-labelled 30 residues long poly-L-lysine. Labeling of the parasite membranes with a distinct enrichment in the middle of the sporozoites is noticeable. Images are single representation of the multiple incubated *P. falciparum* sporozoites. Each channel for microscopy is indicated and bar represents 4.2 μM. Imaging was done on Zeiss AxioImager M2 fluorescence microscope.
